# Transcranial Magnetic stimulation (TMS) modulates functional activity of SH-SY5Y cells: An in vitro model provides support for assumed excitability changes

**DOI:** 10.1101/2020.08.19.257295

**Authors:** Alix C. Thomson, Tom A. de Graaf, Teresa Schuhmann, Gunter Kenis, Alexander T. Sack, Bart P.F. Rutten

## Abstract

Repetitive Transcranial Magnetic Stimulation (rTMS) is an established neuromodulation technique, using electromagnetic pulses that, depending on the precise parameters, a*re assumed* to lead to lasting neural excitability changes. rTMS has widespread applications in both research and therapy, where it has been FDA approved and is considered a first-line treatment for depression, according to recent North American and European guidelines. However, these assumed excitability effects are often difficult to replicate, and highly unreliable on the single subject/patient level. Given the increasing application of rTMS, especially in clinical practice, the absence of a method to unequivocally determine effects of rTMS on human neuronal excitability is problematic. We have taken a first step in addressing this bottleneck, by administering excitatory and inhibitory rTMS protocols, iTBS and cTBS, to a human *in vitro* neuron model; differentiated SH-SY5Y cells. We use live calcium imaging to assess changes in neural activity following stimulation, through quantifying fluorescence response to chemical depolarization. We found that iTBS and cTBS have opposite effects on fluorescence response; with iTBS increasing and cTBS decreasing response to chemical depolarization. Our results are promising, as they provide a clear demonstration of rTMS after-effects in a living human neuron model. We here present an in-vitro live calcium imaging setup that can be further applied to more complex human neuron models, for developing and evaluating subject/patient-specific brain stimulation protocols.

## 1. Introduction

Transcranial Brain Stimulation describes all forms of neuromodulation in which neural activity is stimulated noninvasively by applying electric or electromagnetic pulses through the intact skull into the brain. In Transcranial Magnetic Stimulation (TMS), magnetic pulses are applied transcranially to induce action potentials (Barker, Jalinous, & Freeston, 1985). When multiple TMS pulses are administered in a particular pattern/frequency (repetitive, rTMS) long-lasting effects on cortical excitability have been described (Huang, Edwards, Rounis, Bhatia, & Rothwell, 2005; Pascual-Leone, Valls-Sole, Wassermann, & Hallett, 1994). Depending on parameters, a targeted cortical brain region can either show lasting increases or decreases of cortical excitability, having widespread implications both in research and the clinic. rTMS has proven effective as a treatment for various mental disorders; such as treatment resistant depressive disorder (George et al., 1995), offering a cost-efficient, painless alternative to pharmaceutical treatment with minimal side effects and risk (O’Reardon et al., 2007; Voigt, Carpenter, & Leuchter, 2017). However, to fully deliver on this potential, the modulatory effects of rTMS protocols must be understood, reliable, and ideally patient-tailored.

Theta Burst Stimulation is the prime example of a rapid rTMS protocol in which opposing neuroplastic effects can be induced depending on the chosen frequency/pattern of stimulation. “Theta burst” refers to triplet bursts (50 Hz) of magnetic stimulation administered at theta frequency (5 Hz). It has been shown that *intermittent* Theta Burst Stimulation (iTBS) increases, whereas *continuous* Theta Burst Stimulation (cTBS) decreases cortical excitability, for up to 1 hour following stimulation (Huang et al., 2005). However, recent reports have emphasized the difficulty of replicating the effects of these different TBS protocols (Lopez-Alonso, Cheeran, Rio-Rodriguez, & Fernandez-Del-Olmo, 2014; Rocchi et al., 2018; Schilberg, Schuhmann, & Sack, 2017; Thomson et al., 2019), raising questions about their robustness and replicability, and subsequently about the generally assumed cellular basis of TBS-induced neuroplastic changes.

Currently in humans, cortical excitability and its modulation by rTMS are almost exclusively measured through motor-evoked potentials (MEP’s), a contralateral muscle twitch which represents cortico-spinal excitability at the time of stimulation (Rothwell et al., 1999). However, this approach is hindered by large inter and intra subject variability (Lopez-Alonso et al., 2014; Schilberg et al., 2017). Alternatively, rTMS-induced changes in cortical excitability can be assessed with EEG; for example with TMS-Evoked Potentials (TEP’s) or resting state neuronal power/synchronization. However substantial confounds exist with this method as well (Conde et al., 2019). This lack of a clear, foundational ground truth about the effects of rTMS/TBS on human neurons is a problem, as the assumption that specific rTMS protocols alter excitability in a specific direction is the basis for the current and increasing application of rTMS protocols around the world. Perhaps the current inability to reliably verify rTMS effects could be solved by moving from human *in vivo* to human *in vitro* models.

Here, we took a first step by developing a system of functional imaging in SH-SY5Y human neuroblastoma cells, which we used as a model for neuronal activity and excitability following rTMS protocols. SH-SY5Y cells can be relatively quickly and consistently differentiated into a mature neuron-like state, developing functional synapses and expressing many markers of mature human neurons (Jahn et al., 2017; Kaplan, Matsumoto, Lucarelli, & Thiele, 1993; Krishna et al., 2014; Pezzini et al., 2017). We therefore opted to first test our setup in these cells, with the aim of moving to a more complex human neuronal in vitro setup in the future.

We used calcium imaging to measure changes in neuron *activity* following cTBS and iTBS stimulation, and, crucially, to model effects on *excitability* through response to chemical depolarization. To measure calcium activity, we used Fluo-4 AM (F14201, Thermo Fisher), a fluorescent indicator which binds intracellular calcium, to quantify changes in calcium concentration in the 100nM-1mM range (Gee et al., 2000). In a neuron, the resting calcium levels range between 50-100nM, which can increase 100-fold during electrical activity (Berridge, 1998). Therefore, we expected to measure low fluorescent signal at baseline, and an increase in signal intensity with cellular activity. To assess effects *excitability*, or ‘evoked’ functional activity, we measured responses to exposure of 1M Potassium Chloride (KCl), which has been shown to immediately induce cellular activity during calcium imaging in differentiated SH-SY5Y cells (Jahn et al., 2017).

Mature SH-SY5Y neurons were incubated with Fluo-4,AM dye, and imaged for baseline activity with fluorescence microscopy. We then stimulated our human neuron model with commonly used stimulation protocols: iTBS, cTBS, or sham at an intensity of 100% Maximum Stimulator Output (MSO) and a distance to 1cm from the dish surface. This has been done previously in *in vitro* and animal studies of rTMS effects (Hellmann et al., 2012; B. Li et al., 2017). The stimulation setup can be seen in Figure 1A. The distribution of the electric field (V/m) induced by rTMS within the cell culture dish from several viewpoints can be seen in figure 1B-D, modelled with SimNIBS (Thielscher, Antunes, & Saturnino, 2015) using the cell culture dish mesh generously shared by the authors of (Lenz et al., 2016). Post stimulation activity was the measured, followed by the addition of 1M KCl to induce cellular activity. A time-series montage showing an example of the increase in fluorescence in 1 cell as the KCl is added can be seen in in Figure 1E. We measured fluorescence intensity in 4 separate 2-minute blocks; 1. Baseline, 2. Post-stimulation, 3. During the addition of 1M KCl to induce depolarization, and 4. Post-KCl.

**Figure 1.**
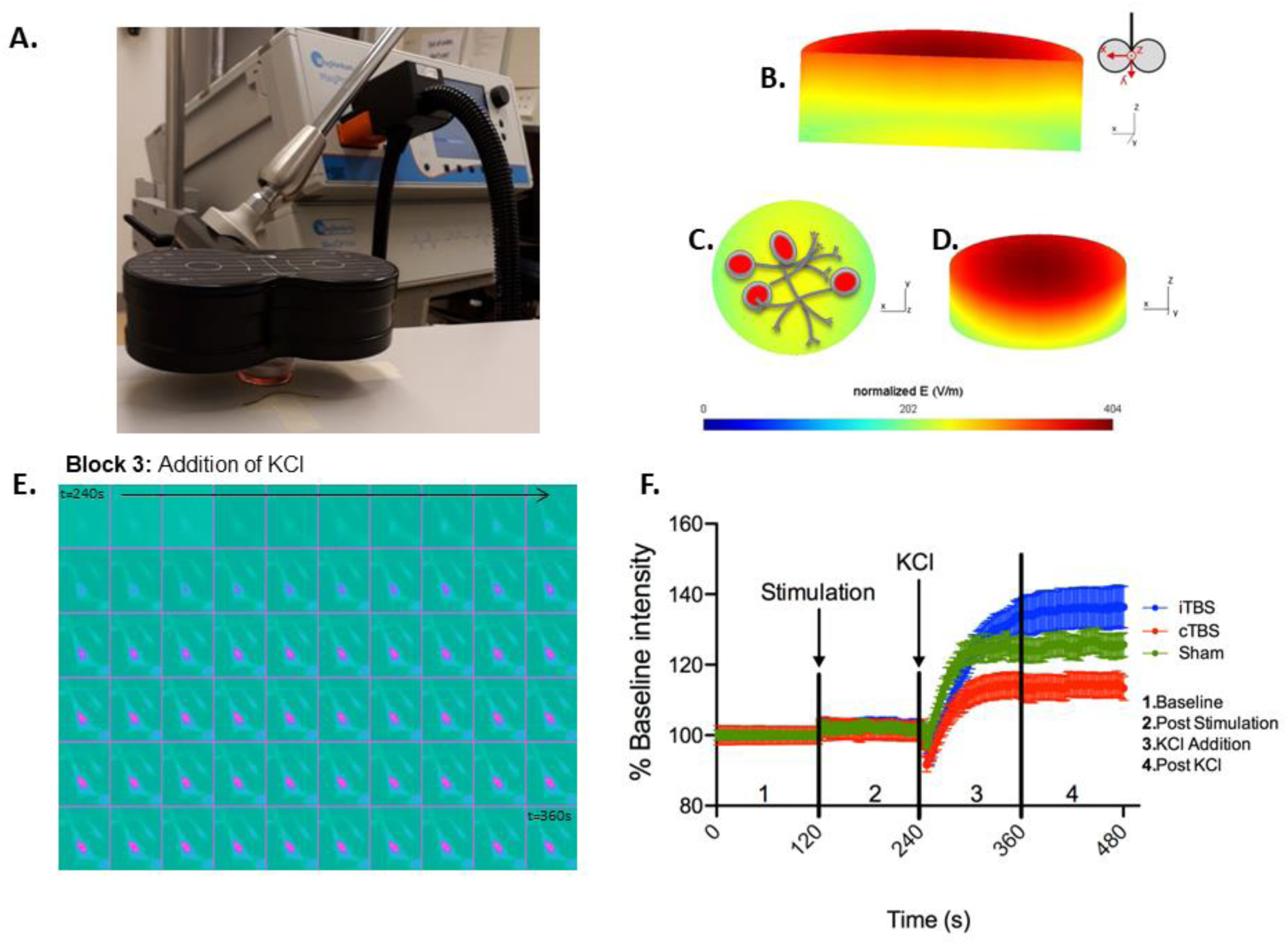
**A.** Position of the cell culture dish 1cm below the center of the coil. **B-D**. Simulation of the induced electric field (V/m) within the cell culture dish. SimNIBS (Thielscher et al., 2015) was used to calculate the electric field induced within the cell culture dish, during TMS stimulation at 100% MSO. The simulation parameters (cell culture dish model and conductivity values) were generously shared by (Lenz et al., 2016). **B**. A cross section of the cell culture dish, showing the gradient of induced electric field within the dish. The electric field is strongest (red) at the top of the dish, closest to the coil. Coil orientation shown beside. **C**. The bottom surface of the cell culture dish (furthest away from the coil, where the cells are plated) and **D**. Tilted view of the dish from the top surface. **E**. Example live cell fluorescence response to 1M KCl stimulation. Each of the 60 squares is a picture of the cell at 2 second intervals, for a period of 2 minutes as the KCl is added. Pink indicates an increase in fluorescence. **F**. Change in fluorescence intensity over time. Each line represents a different cell culture dish. Each data point is the mean % baseline intensity of 20 cells, error bars are standard error of the mean. Block 1 (0-120s): Baseline measurement. After this, cells were removed from the microscope and stimulated with iTBS, cTBS or sham, and placed back into the microscope. Block 2 (120-240s): Post Stimulation measurement. Block 3 (240s-360s): Addition of 1M KCl during recording. Block 4 (360-480s): Post KCl addition.

## 2. Materials and Methods

### Experimental model

SH-SY5Y (ATCC® CRL2266™) neuroblastoma cell line, were used. Freezing and thawing of cell batches were performed according the provided protocols. Cells were not used above passage number 26. For all experiments, cells were grown in DMEM/Nut Mix F12 with Glut-L (Gibco^™^, Thermo Scientific,31331-028) at 37°C and 5% CO^2^. For maintenance and expansion of cell cultures, media was supplemented with 10% heat inactivated Fetal Bovine Serum (FBS, MERCK), 1% penicillin-streptomycin (P/S) and 1% L-Glutamate.

For differentiation, cells were plated in round 35mm Poly-D-Lysine coated 10mm diameter glass bottom MatTek dishes (Matex Corp., P35GC-0-10-C) at approximately 2.4×10^4^ cells per well. FBS supplementation was decreased to 3% FBS 3 days prior to the addition of 10□M Retinoic Acid (RA; Sigma-Aldrich, R2625). During differentiation, RA-supplemented media was replaced every 2 days for 10 days. This differentiation protocol has been shown to establish a mature neuron-like phenotype (Encinas et al., 2000). While it has been reported that the addition of BDNF can promote differentiation and cell survival (Encinas et al., 2000), we did not find this method to be superior to the addition of RA alone (data not shown).

### Functional Imaging

For each condition, 6 separate cell dishes were measured. Cells were incubated in 5µM cell permeant calcium indicator Fluo-4 AM (F14201, Thermo Fisher) and 0.02% Pluronic Acid (ThermoFisher,P6867) made in RA-supplemented culture media for 20 minutes at 37°C and 5% CO_2_. Cells were washed 3 times for 5 minutes in RA-supplemented culture media, the first wash containing 0.00002% Propidium Iodide (Molecular Probes, P3566) and 0.00001% Hoechst (Sigma, B2261). After washing, 1ml of Hank’s Balanced Salt Solution (HBSS) was added to each dish for imaging.

Cells were imaged with the 10X magnification lens of an Olympus IX81 microscope (Olympus Nederland B.V), EXi Blue Fluorescence Camera (Q Imaging, Canada), and X-Cite 120 series fluorescent illuminator (EXFO Photonic Solutions Inc, Canada). A Pecon Tempcontrol 37-2 Digital 2-Channel heating plate (Meyer Instruments, Houston) was used to ensure cell dishes were kept at 37°C during imaging. Imaging was recorded using micromanager open source software (Edelstein et al., 2014). To visualize the change in calcium response over time, 60 images with 2 seconds between each image were recorded per recording block: Baseline, Post Stimulation, KCl Addition and Post KCl. Microscope fluorescence settings are listed in Supplementary Table 1. Following baseline recording, cells were removed from the microscope chamber and stimulated with rTMS as described below. Dishes were marked before removal in order to place them back in approximately the same position.

### Magnetic Stimulation

Cells were placed 1 cm below the centre of a Cool-B65 figure 8 coil (Magventure, Denmark) and stimulated at 100% Maximum Stimulator Output (MSO) with a MagPro X100 with MagOption stimulator (Magventure, Denmark). Each stimulation session consisted of the Huang (2005) published protocol of 50Hz triplets repeated at 5 Hz. iTBS consisted of 2000ms trains with 8000ms inter-train intervals, while cTBS consisted of continuous triplets; both for 600 pulses (Huang et al., 2005). For the cTBS condition, cells were placed under the coil for an additional 150 seconds to ensure iTBS and cTBS cells were out of the incubator for the same amount of time. For the sham condition, cells were placed under the coil for 190 seconds. The field induced within the dish was calculated using SimNIBS toolbox (Thielscher et al., 2015). Electrical conductivities used in the simulation were the same as in Lenz et al., (2016), and the electric dish model used for modelling was generously shared by the authors (Lenz et al., 2016). The rate of change of the coil current was 143 A/μs, corresponding to an intensity of 100% MSO. The resulting normalized electric field induced within the dish from several viewpoints can be seen in Figure 1B-D.

### KCl addition

KCl was added drop by drop into the dish through a needle attached to a large 200ml plastic syringe, to reach a final KCl concentration in the dish of 1 M. This caused an approximately 200-fold increase in the extracellular potassium concentration. This high concentration of KCl required to visualize calcium activity in SH-SY5Y cells has been described previously in the literature (Jahn et al., 2017). The 1 M KCl was left on the cell cultures, and a multi-channel image was then taken to quantify ROI’s after KCl and to visualize cell morphological response to high KCl concentration, followed by another video to quantify stable KCl response for 2 minutes following addition. Fluorescence levels remained high in this measurement block, as has been shown previously in SH-SY5Y cells (Jahn et al., 2017).

### Quantification of Functional Imaging

Fiji (ImageJ, version 1.52i, RRID:SCR_002285) open source software (Schindelin et al., 2012) was used to analyze microscope-acquired images. Each condition (iTBS, cTBS, or sham), was measured from a different cell culture dish. The data is separated into 4 measurement blocks; 1. Baseline, 2. Post-stimulation, 3. During the addition of 1M KCl, and 4. Post-KCl.

First, 20 circular ROI’s (width: 25, height: 25 pixels) of responding cells were randomly selected from the 4^th^ (Post KCl) block, and used to measure pixel intensity over time for every prior block of the measurement. Since the cell culture dish was removed between the baseline and post-stimulation blocks, these ROI’s were shifted between the blocks and if possible, they were adjusted manually to quantify the same cell. Analysis was performed blinded to the condition. Measured intensity values were exported to excel and Prism 5 (Graphpad Software, USA, RRID:SCR_002798.) for statistics and graphing.

Intensity values were normalized to the average of the baseline intensity at each time point. All 20 ROI’s in the baseline block were averaged, to calculate a baseline average at each time point (BaseAv(T_x_)). Each intensity value for each ROI was then divided by the average baseline intensity at that time point (Intensity (T_x_)/BaseAv(T_x_))*100 to give a percent change from baseline.

The mean % baseline intensity of 20 ROI’s for each time point, with error bars as standard error of the mean can be seen in Figure 1F. ROI outliers which were 2.5 standard deviations above the mean were removed. This resulted in the removal of 16 ROIs of the total 120.

### Statistical Analysis

Analysis was done in Prism 5 (Graphpad Software, USA, RRID:SCR_002798) combining data from all 6 independent experiments. 2-way ANOVA with Bonferroni-corrected post-hoc comparisons were used to test the effect of CONDITION (iTBS, cTBS, sham) and TIME (60 time point measurements per block) on % baseline fluorescence intensity in each of the measurements blocks; 1.Baseline 2. Post-stimulation, 3. Addition of KCl, 4. Post-KCl.

## 3. Results

We found that neurons which had been stimulated with iTBS, a protocol assumed to increase excitability, showed greater increase in fluorescence response to KCl than those which had been sham stimulated. In striking contrast, neurons which had been stimulated with cTBS, a protocol assumed to decrease excitability, showed the opposite response to KCl, i.e. a decreased response. A representative plot of the mean % change from baseline fluorescence during the entire experiment (error bars are standard error of the mean) is shown in Figure 1F.

We repeated this experiment in 6 independent cell cultures per stimulation condition (iTBS, cTBS, or sham). For statistical analysis, data from all experiments were combined. The mean % baseline intensity of 20 cells per experiment was calculated for each time point, for each experiment, to give an n=6, each with 60 time points per block. On the group level in block 2 (Post Stimulation), a two way ANOVA with factors condition (iTBS, cTBS, sham) and time reveals a significant effect of condition (F(2,660)=83.53, p<0.0001), and no effect of time (F(59,660)=0.01716, p=1.00) or interaction (F(118,660)=0.01366, p=1.000). Post-hoc Bonferroni corrected planned comparisons to % baseline intensity after sham stimulation (101.4±0.23%) show a minor but significant decrease (p<0.0001) in baseline intensity after cTBS (99.27±0.08%), but no significant increase (p=0.1287) after iTBS (102.0±0.23%). Thus, cTBS/iTBS had very small, though significant, effects on ‘baseline’ neuronal activity as captured by pre-stimulation normalized fluorescence.

As hypothesized, TBS modulations were particularly apparent on ‘evoked’ neuronal activity. This functional activity was captured by the neuronal response to chemical depolarization (KCl), in block 4. We found a greater increase in fluorescence response to KCl in neurons stimulated with iTBS (125.4±0.44%) as compared to sham (122.2±0.42%). Neurons stimulated with cTBS instead showed a decreased fluorescence response to KCl (114.8±0.65%). A two-way ANOVA of baseline-corrected intensity following KCl addition confirms a statistically significant, strong effect of condition (F(2,840)=91.46, p<0.001), but no effect of time (F(59,840)=0.04, p=1.000) and no interaction (F(118,840)=0.035, p=1.000). iTBS and cTBS comparisons to sham stimulated neurons showed statistically significant differences (p<0.001, Bonferroni-corrected). See Supplementary Information for 2-way ANOVA results of each block. Figure 2 shows the mean % baseline intensity for each block in each stimulation condition.

**Figure 2.**
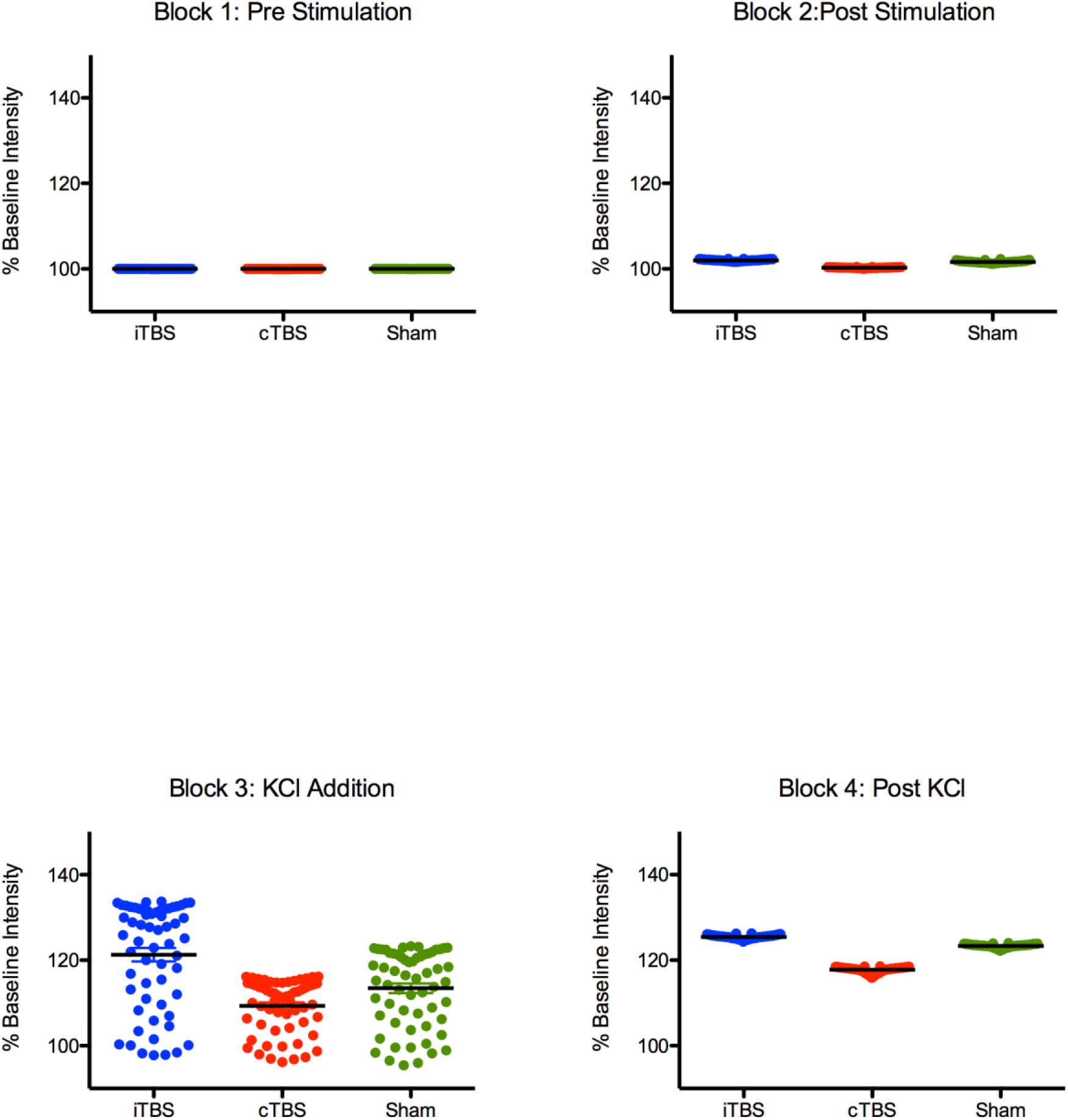
Group Results of each stimulation condition, for each block. Each dot is the mean time point intensity of 6 independent experiments. 120 time points are combined in each block; the black line is the mean % baseline.

## 4. Discussion

Taken together, these findings provide direct evidence that rTMS was able to modulate neuronal response to chemical depolarization with KCl; with iTBS facilitating, and cTBS inhibiting this response, compared to sham stimulation. This provides direct and new support for the hypothesized cTBS and iTBS effects on human neuronal excitability.

These findings, providing support for the positive and negative modulation of human neuronal excitability by iTBS and cTBS respectively, are reassuring. While they do not explain reported unreliability of TBS effects on other human excitability measures (e.g. MEP/TEP), they 1) validate at least on the neuronal level foundational assumptions about the mechanistic underpinnings of these globally implemented treatment/research protocols, and 2) demonstrate the value of the presented *in vitro* human neuron model for studying cellular effects of TMS (and potentially for other interventions and exposures linked to the onset or treatment of mental disorders). Therefore, this functional imaging setup of stimulated living human neurons in a dish provides a direct method for testing the immediate functional activity changes of human neurons following TMS.

Animal studies have previously provided support for the hypothesized opposing neuroplastic effects of iTBS and cTBS (Benali et al., 2011; Volz, Benali, Mix, Neubacher, & Funke, 2013), as well as an immediate effect of rTMS on intracellular calcium release (Banerjee, Sorrell, Celnik, & Pelled, 2017; Grehl et al., 2015). Computational modelling of calcium-dependent plasticity effects following rTMS in a neural field model have shown that during iTBS/cTBS stimulation, the first burst of stimulation within a train causes an increase in calcium concentration, resulting in potentiation (Wilson, Fung, Robinson, Shemmell, & Reynolds, 2016). In an iTBS protocol, 8 second breaks between trains of stimulation bursts allow for the maintenance of this potentiating increase in calcium concentration. In cTBS, the accumulation of calcium concentration results in an overall depressive effect (Wilson et al., 2016). Previous studies have also discussed the importance of the intervals between stimulation trains in the iTBS protocols for it’s faciliatory effect (Hamada, Murase, Hasan, Balaratnam, & Rothwell, 2013). In line with these findings, we also report an immediate depressive effect of cTBS and an immediate faciliatory effect of iTBS on calcium concentration, as measured by fluorescence microscopy.

We modelled the induced electric field within the cell culture dish using SimNIBS as described in the methods. With stimulation at 100% MSO and 1cm above the dish, which is stronger than what is typically used for human cortex stimulation. In our cell culture dish, all neurons were affected by an electric field greater than 200V/m, while in models of the human cortex effected by stimulation, only a very superficial layer of cortex directly under the coil is effected by an electrical field of approximately 100V/m (Deng et al., 2019; Lu & Lu, 2018). However, other rTMS animal and cell culture studies have reported using these stimulation parameters (Hellmann et al., 2012; B. Li et al., 2017), and in this first, exploratory setup we wanted to measure changes in response to the highest stimulation intensity possible.

One of the main limitations of this study, is the use of SH-SY5Y cells as a model for human neurons. SH-SY5Y cells are a human-derived neuroblastoma cell line, which can be differentiated to neural-like cells in a relatively short amount of time (Encinas et al., 2000; Kovalevich & Langford, 2013; Shipley, Mangold, & Szpara, 2016). They are well characterized, have been shown to develop functional synapses (Jahn et al., 2017; Toselli, Tosetti, & Taglietti, 1996), synthesize neurotransmitters (Krishna et al., 2014), and are used as a model for various human neuropsychiatric and neurodegenerative disorders (Agholme, Lindstrom, Kagedal, Marcusson, & Hallbeck, 2010; Henkel et al., 2008; Xicoy, Wieringa, & Martens, 2017). However, they are not glutamatergic or GABAergic neurons, which are thought to be the primary cortical neurons affected by rTMS (Cirillo et al., 2017). SH-SY5Y cells have been shown to express many genes involved in axonal guidance and synaptic plasticity (Agholme et al., 2010; Jahn et al., 2017; Kaplan et al., 1993; Pezzini et al., 2017), specifically the BDNF-TrkB pathways, known to be involved in LTP (Minichiello et al., 1999). Therefore, while not the perfect human neuron model, we chose to first test our setup with SH-SY5Y cells as they are quick to culture, resulting in relatively reliable and stable phenotypes and they have been used as a model for human synaptic plasticity *in vitro*.

Recently, brain-state has been suggested as a factor contributing to variability of individual responses to rTMS protocols (Bergmann, 2018). rTMS is thought to lead to LTP/LTD-like synaptic plasticity effects, however, on a meta-plastic level homeostatic mechanisms work to stabilize neural systems, keeping the threshold at which synaptic plasticity is induced within a physiologically relevant range (Karabanov et al., 2015; J. Li, Park, Zhong, & Chen, 2019; Turrigiano, 2008; Turrigiano & Nelson, 2004). These metaplasticity mechanisms are thought to adjust this threshold for synaptic plasticity based on previous neural activity (Abraham & Bear, 1996; Bienenstock, Cooper, & Munro, 1982), therefore brain-state is extremely influential in determining the excitability effects induced by rTMS. In our cell culture model, we did not test for the influence of metaplastic or homeostatic plasticity mechanisms, as we were interested in 1.) establishing an *in vitro* setup capable of measuring excitability changes following rTMS in a human neuron model, and 2.) using this setup to quantify the *immediate* excitability effects of iTBS and cTBS. Additionally, we have not yet proven that the functional activity effects we find are related to synaptic plasticity, which would be a necessary next step in establishing the presumed meta-or homeostatic plasticity effects of rTMS. Based on our work, future studies can be designed to specifically test the effects history of neural activity on TMS-induced plasticity changes in a human neural network model.

We here present a setup for measuring changes in human neuron activity following rTMS in living human neurons. This method is feasible, reliable, and can be used to develop and test existing, new, optimized and individualized rTMS protocols for various applications. Looking ahead, this human neuron model has the potential to allow laboratory evaluation and optimization of rTMS protocols for individual patients, particularly when using patient-derived cells (e.g. skin-derived fibroblasts or peripheral blood mononuclear cells) and induced pluripotent stem cell (iPSC) reprogramming (Takahashi & Yamanaka, 2006). For example, using iPSC techniques, individualized neuron models can be established by reprogramming skin cells taken from a treatment resistant patient into neurons which contain the genetic composition of that patient (Ambasudhan et al., 2011; Takahashi & Yamanaka, 2006). This can be taken one-step further with the cerebral organoid, as these patient specific neurons can organize into a 3D cell culture system to model whole cortical structures *in-vitro* (Lancaster et al., 2013). With this setup, patient-specific human neural models can be stimulated with rTMS, using calcium imaging to measure that patient’s responsiveness to particular rTMS protocols. Further application and development of this model may enable assessment and exploration of planned patient-tailored rTMS treatments, thus optimising patient care.

## Acknowledgements

Funding Source: This work was supported by the Netherlands Organization for Scientific Research (NWO, Veni to T.A.G 451-13-024; Vici to A.T.S 453-15-008), and an internal grant from the Centre for Integrative Neuroscience at Maastricht University.

## CRediT

**Alix Thomson:** Conceptualization, Methodology, Formal analysis, Investigation, Writing-Original Draft. **Gunter Kenis:** Conceptualization, Writing-Review & Editing, Supervision, Project administration. **Tom A. de Graaf:** Writing-Review & Editing, Supervision. **Teresa Schuhmann:** Writing-Review & Editing, Supervision. **Alexander T. Sack:** Writing-Review & Editing, Supervision, Funding Acquisition. **Bart P.F. Rutten** Writing-Review & Editing, Supervision, Funding Acquisition.

## Conflict of Interest Statement

None of the authors have potential conflicts of interest to be disclosed.

## Notes

### Competing Interest Statement

The authors have declared no competing interest.

